# Tonic sound-evoked motility of cochlear outer hair cells in mice with impaired mechanotransduction

**DOI:** 10.1101/2024.12.19.629412

**Authors:** James B. Dewey

## Abstract

Cochlear outer hair cells (OHCs) transduce sound-induced vibrations of their stereociliary bundles into receptor potentials that drive changes in cell length. While fast, phasic OHC length changes are thought to underlie an amplification process required for sensitive hearing, OHCs also exhibit large tonic length changes. The origins and functional significance of this tonic motility are unclear. Here, *in vivo* cochlear vibration measurements reveal tonic, sound-induced OHC motility in mice with stereociliary defects that impair mechanotransduction and eliminate cochlear amplification. Tonic motility in impaired mice was physiologically vulnerable but weakly related to any residual phasic motility, possibly suggesting a dissociation between the underlying mechanisms. Nevertheless, a simple model demonstrates how tonic responses in both normal and impaired mice can result from asymmetric mechanotransduction currents and be large even when phasic motility is undetectable. Tonic OHC responses are therefore not unique to sensitive ears, though their potential functional role remains uncertain.

## INTRODUCTION

The sensitivity and frequency selectivity of mammalian hearing depends on a mechanical amplification process mediated by cochlear outer hair cells (OHCs)^1,2^. OHCs detect sound-evoked vibrations of the underlying basilar membrane and organ of Corti structures through deflection of their stereociliary bundle, leading to current flow through mechanotransduction channels located at the stereocilia tips (**Fig. 1a**). The resulting changes in membrane potential drive conformational changes in prestin, a motor protein densely expressed in the OHC lateral wall, causing the OHCs to change length and generate force^3–6^. This electromotile force generation is commonly thought to operate fast enough to follow changes in membrane potential at acoustic frequencies, thus amplifying organ of Corti vibrations on a cycle-by-cycle basis^7–9^. However, in addition to phasic, cycle-by-cycle motility, OHCs also exhibit slow or tonic length changes in response to a variety of electrical, chemical, and mechanical stimuli *in vitro*^3,10–13^, as well as in response to sound *in vivo*^9,14–18^. While slow or tonic OHC length changes have been proposed to regulate cycle-by-cycle electromotility or otherwise modulate cochlear vibrations^18,19^, their precise origin and functional role, if any, remain poorly understood.

**Figure 1.**
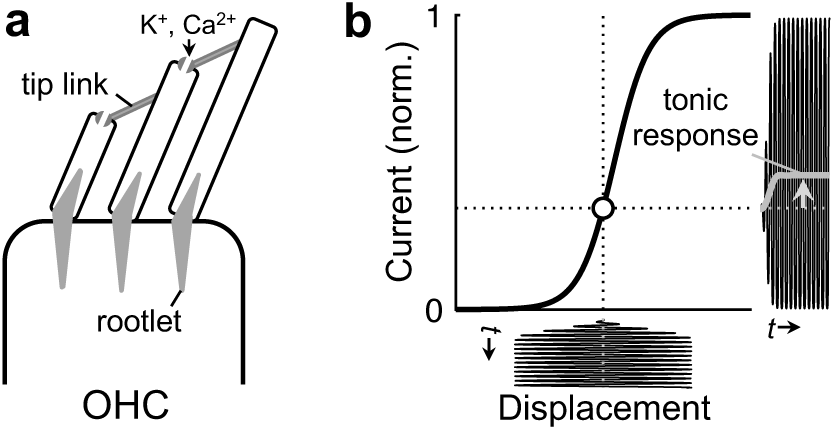
Theoretical dependence of tonic OHC responses on asymmetric mechanotransduction. (**a**) Schematic of the top of an OHC and its stereociliary bundle. The stereocilia are anchored to the apical OHC surface by stiff rootlets. Deflection of the bundle stretches the tip links that connect adjacent stereociliary rows, thus opening ion channels. (**b**) First-order Boltzmann function approximating the relationship between bundle displacement and the resulting mechanotransduction current. If the bundle’s resting position is biased away from the center of the function, sinusoidal bundle displacements will produce asymmetric currents that contain a tonic (i.e., direct current or 0 Hz) component. Low-pass filtering the current waveform allows direct visualization of the tonic component (gray curve).

*In vivo*, sound-evoked tonic OHC length changes could theoretically be driven by sustained changes in membrane potential. Such changes would naturally arise if OHC mechanotransduction currents were asymmetric, resulting in a net depolarization or hyperpolarization during stimulation (**Fig. 1b**). This would lead to a tonic OHC contraction or elongation, respectively. While OHC mechanotransduction currents have often been proposed to be roughly symmetric^20,21^, direct mechanical measurements and intra- or extracellular potentials from several species have revealed significant tonic components and related even-order distortions that are indicative of asymmetric nonlinearity^9,16,17,22,23^. Sound-evoked tonic OHC movements depend on normal prestin function and largely disappear postmortem^9^, supporting the interpretation that these motions are due to the electromotile response to a mechanotransduction-induced change in membrane potential. Nevertheless, the dependence of these tonic OHC responses on the integrity and nonlinearity of the mechanotransduction process has yet to be directly demonstrated.

Here, this relationship was examined by measuring sound-evoked cochlear vibrations in mice with mutations specifically affecting the stereociliary bundle. These included *salsa* mice, which progressively lose the stereociliary tip links that are thought to gate mechanotransduction channels^24^, and *Triobp*^Δex8/Δex8^ mice, which lack stereociliary rootlets, resulting in bundles that are less stiff and prone to damage, such that they degenerate by adulthood^25^. In both mice, OHC-mediated amplification is severely impaired, presumably due to a loss of mechanotransduction^26^. Surprisingly, tonic OHC displacements were observed in a significant fraction of these impaired mice. While the results could indicate a potential distinction between the mechanisms underlying tonic and phasic OHC motility, a simple model of OHC responses demonstrates how realistic changes to the mechanotransducer function can result in the persistence of tonic motility despite the apparent absence of cycle-by-cycle OHC feedback. The findings therefore highlight a form of nonlinear OHC activity that may remain in ears with certain types of hearing loss.

## RESULTS

### Sound elicits tonic OHC motility in wild-type mice with normal cochlear amplification

Optical coherence tomography (OCT) was used to image the mouse cochlear apex and measure sound-evoked vibrations from within the organ of Corti (**Fig. 2a-c**). Responses were first characterized in wild-type (WT) C57BL/6J mice, as the impaired mouse strains were both on this background. In each mouse, the presence of cochlear amplification was initially assessed by measuring the phasic displacements of the basilar membrane (BM) in response to tones varied both in frequency and sound pressure level (SPL).

**Figure 2.**
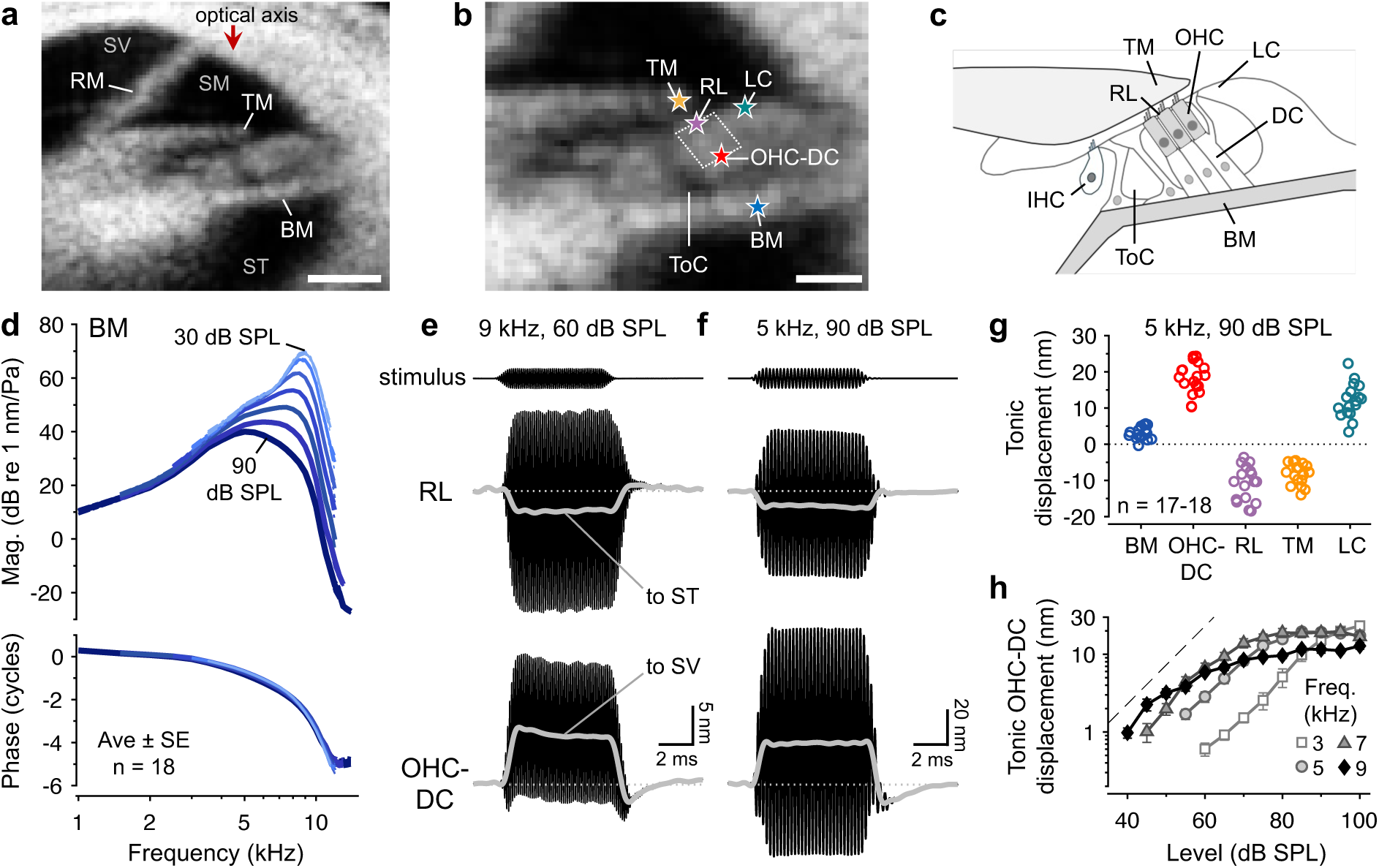
Sound evokes tonic OHC contractions in the apex of the WT mouse cochlea. (**a**) Cross- sectional OCT image of an apical cochlear location tuned to ∼9 kHz in a WT C57BL/6J mouse. The three fluid-filled scalae (SV = scala vestibuli, SM = scala media, ST = scala tympani) are indicated along with Reissner’s membrane (RM), the tectorial membrane (TM), and basilar membrane (BM). Scale bar = 100 µm. (**b**) Magnified view of the organ of Corti indicating the approximate location of the OHCs (dotted white lines), which was inferred from the positions of the tunnel of Corti (ToC) and TM. Stars indicate measurement points where tonic displacements were measured (RL = reticular lamina, LC = lateral cell region, OHC-DC = OHC-Deiters’ cell junction). Scale bar = 50 µm. **(c)** Organ of Corti schematic with relevant anatomy labeled (IHC = inner hair cell, DC = Deiters’ cell). (**d**) Average BM displacement magnitudes and phases as a function of stimulus frequency and sound pressure level (SPL) in WT mice. Displacement magnitudes are normalized to the evoking stimulus pressure (in Pascals, Pa) to highlight the nonlinearity exhibited by the responses. Responses to 10 and 20 dB SPL tones were also obtained but are not shown, for clarity. Dashed-dotted lines indicate ± 1 SE, though these are generally obscured by the average curves. (**e-f**) Displacement waveforms measured from the RL and OHC-DC junction for a 9 kHz, 60 dB SPL stimulus (**e**) and a 5 kHz, 90 dB SPL stimulus (**f**) in an individual mouse. Low-pass filtered waveforms (gray curves) reveal that the RL and OHC-DC junction tonically moved toward one another during the stimulus. Different displacement scales are used in **e** and **f**, and stimulus pressure waveforms are arbitrarily scaled. (**g**) Tonic displacements measured from the OHC region and surrounding structures for a 5 kHz, 90 dB SPL stimulus in WT mice (n = 18 for the OHC-DC junction; n = 17 for all other points). Positive and negative displacements are toward SV and ST, respectively, and are plotted on a linear scale. (**h**) Average tonic displacements of the OHC-DC junction vs. stimulus level at four frequencies (n = 5, 13, 6, and 6 mice for 3, 5, 7, and 9 kHz stimuli, respectively). Displacements are plotted on a logarithmic scale and the dashed line illustrates linear growth. Error bars indicate ± 1 SE.

In 3-7 week-old WT mice, BM responses were tuned to a characteristic frequency (CF) of ∼9 kHz and exhibited the frequency- and level-dependent nonlinearity that is a hallmark of cochlear amplification (**Fig. 2d**). To highlight the nonlinearity, average displacement magnitudes are shown normalized to the stimulus pressure, yielding the “gain” of the responses relative to the stimulus. For frequencies near the CF, the gain of BM responses to low-level stimuli was ∼40 dB higher than the gain for high-level stimuli. This nonlinear gain is attributed to the saturation of OHC mechanotransduction currents with increasing stimulus level, which leads to the compressive growth of the receptor potential and the OHC’s electromotile response. At lower frequencies, BM response gains became more similar across stimulus level, indicating linear growth. Less level dependence was observed for the phases of BM displacements, which showed increasing lags with frequency that result from traveling wave propagation.

After assessing the frequency tuning and nonlinear gain of BM vibrations, responses to short tones were obtained from the bottom and top of the OHC region to characterize any tonic OHC length changes. As shown previously in CBA/CaJ mice, displacement waveforms measured from the OHC region in C57BL/6J mice were asymmetric, revealing motions that were consistent with tonic OHC contraction (**Fig. 2e-f**). Tonic displacements were visualized by low-pass filtering the waveforms (gray curves in **Fig. 2e-f**), which showed that the tops of the OHCs, near the reticular lamina (RL), moved toward scala tympani (ST) while the bottoms of the OHCs, near their junctions with the Deiters’ cells (DCs), moved toward scala vestibuli (SV) for the duration of the stimulus. The structures surrounding the OHCs also exhibited tonic motions that would result from OHC contraction (**Fig. 2g**). The BM moved by a small amount toward SV, consistent with it being a stiff structure that is pulled by the motion of the DCs. In contrast, the TM moved with the RL toward ST, presumably because it is attached to the RL by the tallest row of OHC stereocilia. Lastly, the lateral portion of the organ of Corti’s apical surface, spanning roughly from the third OHC to the Hensen’s cells (here termed the lateral cell, or LC, region) moved toward SV, indicating that this region pivots upward and/or bulges outward when the tops of the more medial OHCs move down toward ST.

Tonic displacements were usually largest at the OHC-DC junction and were reliably elicited by CF tones presented at stimulus levels above 40 dB SPL, as well as by below-CF tones presented at higher levels (**Fig. 2h**). While tonic displacements typically reached at most 10 nm for near-CF stimuli, they could exceed 20 nm at lower frequencies. This is consistent with OHC mechanotransduction being driven by movements that are roughly proportional to BM displacement, which saturates with increasing stimulus level near the CF (reaching ∼20 nm at 90 dB SPL) but continues to grow below the CF (reaching ∼90 nm for a 5 kHz, 90 dB SPL stimulus). This would lead to a larger maximal tonic component in the OHC receptor potential at frequencies below the CF and, thus, larger tonic displacements. At high stimulus levels, tonic displacements peaked near the frequency of maximum BM displacement, which was ∼5 kHz (**Supplementary Fig. S1**).

### Sound elicits tonic OHC motility in a large subset of mice with impaired mechanotransduction

To test the dependence of the tonic displacements on OHC mechanotransduction, responses were measured in 3-7 week-old *salsa* and *Triobp*^Δex8/Δex8^ mice (hereafter referred to as *Triobp* mice). Both mice have stereociliary defects that result in severe deafness by early adulthood^24,25^. BM displacements in *salsa* and *Triobp* mice were tuned to ∼4-5 kHz and generally lacked nonlinear amplification, as indicated by the overlapping displacement gains obtained for 50 to 100 dB SPL stimuli (**Fig. 3a-b**). In two *salsa* mice, modest BM nonlinearity was observed, with displacement gains for 60 dB SPL tones being more than 20 dB higher than those at 100 dB SPL for frequencies near 8-9 kHz. Since this indicates that OHC function was not as dramatically affected in these mice, their responses were not included in any subsequent analyses, nor were their data included in the averages shown in **Fig. 3**. In all other mice, the difference in gains between low and high stimulus levels did not exceed 10 dB, with average values being close to 0.

**Figure 3.**
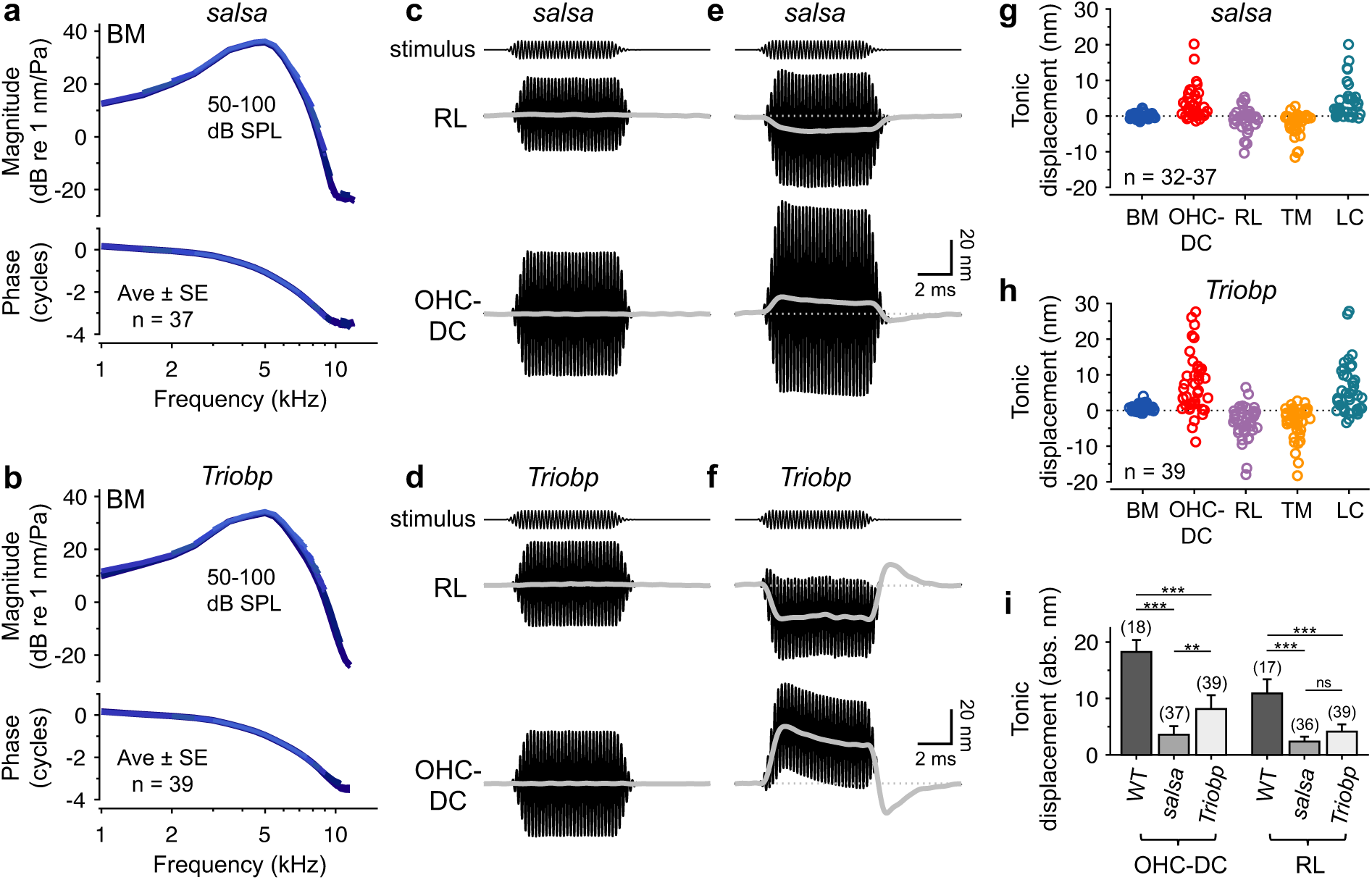
Sound evokes tonic OHC contractions in mice with impaired mechanotransduction and minimal nonlinear BM amplification. (**a-b**) Average BM displacement gain (re stimulus pressure) and phase as a function of stimulus frequency and level for *salsa* (**a**) and *Triobp* (**b**) mice. Displacement gain curves for 50-100 dB SPL stimuli are indistinguishable, indicating the linearity of the responses and the absence of BM amplification. (**c-f**) Displacement waveforms of the RL and OHC-DC junction in individual *salsa* (**c**,**e**) and *Triobp* (**d**,**f**) mice for a 5 kHz, 90 dB SPL stimulus. While tonic displacements could not be detected in some mice (**c-d**), they were present in many others (**e-f**; gray curves are the low-pass filtered waveforms). Stimulus waveforms are arbitrarily scaled. See **Supplementary Fig. S3** for more examples of tonic displacements in impaired mice. (**g-h**) Tonic displacements measured from different points within the OHC region and surrounding structures for 90 dB SPL stimuli in *salsa* (**g**) and *Triobp* (**h**) mice. Positive and negative displacements are toward SV and ST, respectively. Stimulus frequencies were set within 0.5 kHz of the frequency eliciting maximum BM displacement in each mouse and ranged from 4 to 5.5 kHz. (**i**) Average absolute tonic displacement magnitudes for the OHC-DC junction and RL compared across strains (individual data are shown in **g** and **h** and Fig. 2g). The number of mice included in each average is shown in parentheses and error bars indicate 95% confidence intervals. Tonic displacements were compared across strains using one-way ANOVAs (for OHC-DC data, F_2,91_ = 37.58, p < 0.005; for RL data, F_2,89_ = 31.93, p < 0.005). Asterisks indicate statistical significance of post-hoc comparisons with Bonferroni corrections (** p < 0.005, *** p < 0.0005, ns = not significant).

At 7 kHz, the average difference (± standard error, SE) between response gains for 60 and 90 dB SPL stimuli was 0.57 ± 0.19 dB in *salsa* mice (n = 36) and 0.88 ± 0.20 dB in *Triobp* mice (n = 39), compared to 18.73 ± 0.34 dB in WT mice (n = 18).

If tonic OHC responses require normal mechanotransduction, they would be expected to be lost in these impaired mice. However, while tonic displacements were indeed absent in some *salsa* and *Triobp* mice, they could be measured in a surprisingly large subset (**Fig. 3c-f**). For 90 dB SPL tones presented at the frequency eliciting maximum BM vibration, tonic displacements of the OHC-DC junction were greater than the approximate detection threshold of 1.5 nm in 19 of 37 *salsa* mice (51%) and 32 of 39 *Triobp* mice (82%). Tonic displacements in *salsa* mice were generally small, while displacements in *Triobp* could sometimes be as large or larger than those observed in WT mice, approaching 30 nm.

When present, tonic displacements in impaired mice were largest for frequencies eliciting maximal BM displacement (**Supplementary Fig. S2**) and were usually consistent with OHC contraction (**Fig. 3g-h**), as in WT mice. However, in two *Triobp* mice, the RL and OHC-DC junction moved away from each other, indicating elongation. In five *salsa* and three *Triobp* mice, the OHC-DC junction and RL moved in the same direction (toward SV in all but one *Triobp* mouse). Apparent uniform motion of the OHC region could result from the angle of the OHCs relative to the optical axis or imprecision in the measurement points chosen to represent the RL and OHC-DC junction. Alternatively, this may reflect real changes in how tonic OHC forces are coupled to the surrounding structures. Regardless, in most impaired mice with measurable tonic displacements, the motions were similar to those observed in WT mice, albeit with a lower amplitude (**Fig. 3i**).

### Tonic displacements in impaired mice are physiologically vulnerable and reduced with age

Tonic displacements in impaired mice sometimes exceeded the phasic displacement at the stimulus frequency (e.g., see initial portion of the OHC-DC response in **Fig. 3f** and **Supplementary Figs. S3**). This suggests that the tonic displacements did not simply result from OHC region motion being more constrained in one direction versus the other. Instead, the tonic displacements likely reflect the electromotile response to a sustained change in membrane potential, indicating the presence of some form of residual mechanotransduction.

Consistent with this, tonic displacements in all mice were largely eliminated after death, which reduces the endocochlear potential and the strength of any mechanotransduction currents (**Fig. 4a-b**). Tonic displacements obtained 2-45 min postmortem were undetectable in most mice, though response magnitudes of 1.5-5 nm were observed in three WT mice (out of 12 mice), one *salsa* mouse (out of 17), and seven *Triobp* mice (out of 30). While not studied systematically, repeated postmortem measurements in several mice indicated that tonic responses could decrease or increase over time. The origin of such fluctuations is unclear though they may be related to gradual postmortem changes in the endocochlear potential or the mechanical properties of the OHCs and organ of Corti.

**Figure 4.**
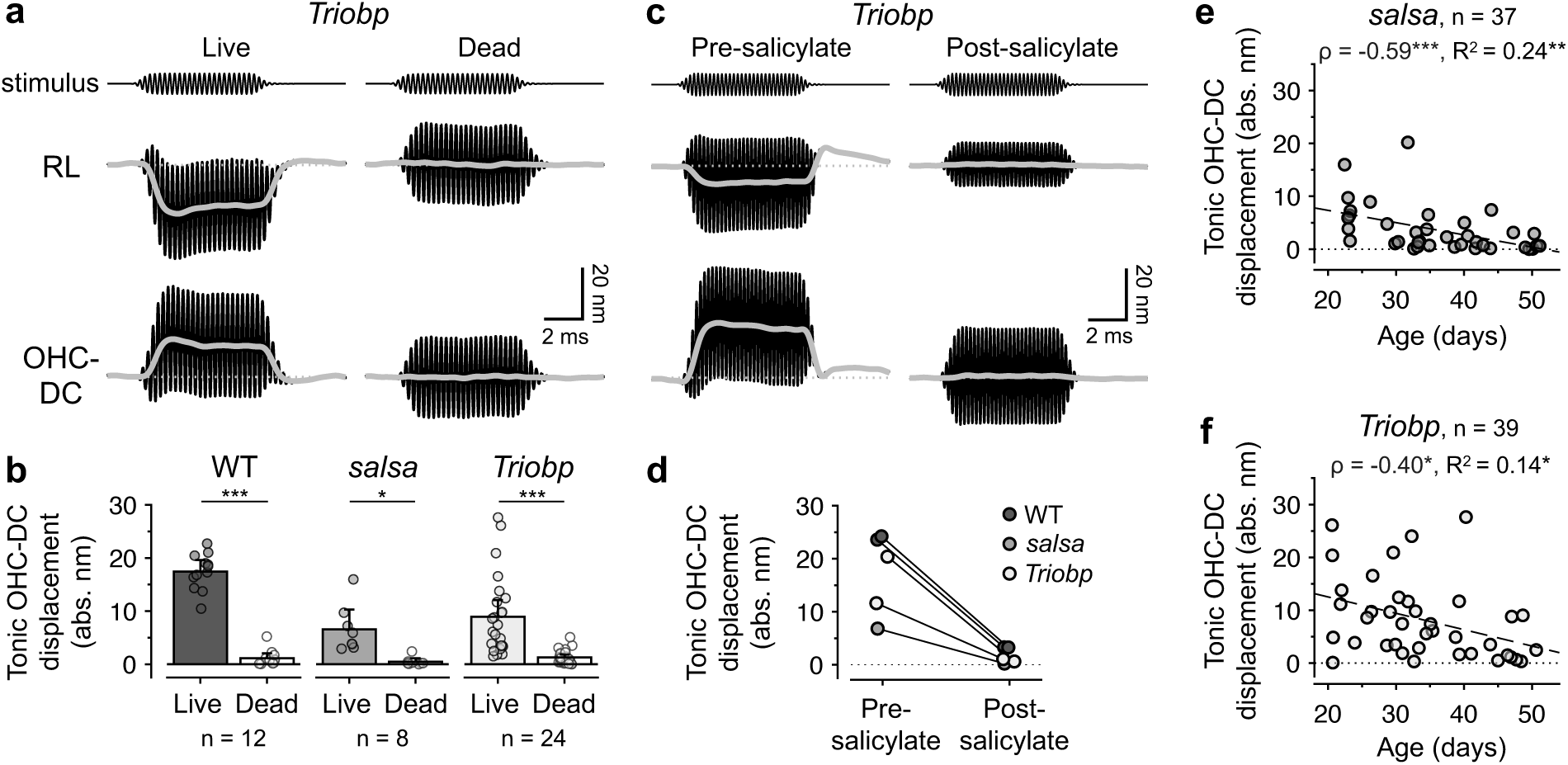
Tonic displacements in impaired mice are diminished after death, salicylate administration, and aging. (**a**) Waveforms of RL and OHC-DC junction displacements for a 4 kHz, 90 dB SPL stimulus in an individual *Triobp* mouse before and ∼40 min after death. Low-pass filtered waveforms (gray curves) show the tonic displacements, which were largely reduced postmortem. (**b**) Average absolute tonic displacements of the OHC-DC junction before and after death in WT, *salsa*, and *Triobp* mice. Averages only include data from mice with absolute tonic displacements ≥ 1.5 nm in the live condition, and for which postmortem measurements were obtained < 45 min after death. Error bars indicate 95% confidence intervals and individual data are shown with semi-transparent symbols. Tonic displacements were significantly reduced postmortem in all strains, as assessed by paired t-test (for WT, t_11_ = 18.38, p < 0.0005; for *salsa*, t_7_ = 3.60, p = 0.009; for *Triobp*, t_23_ = 4.85, p < 0.0005). (**c**) Waveforms of RL and OHC-DC junction displacements for a 5 kHz, 90 dB SPL stimulus in an individual *Triobp* mouse before and ∼20 min after administration of salicylate to the round window membrane. (**d**) Absolute tonic displacements of the OHC-DC junction for a 90 dB SPL stimulus before and < 30 min after salicylate administration in two WT mice, one *salsa* mouse, and two *Triobp* mice (responses in a WT and *salsa* mouse are shown in **Supplementary Fig. S4**). The stimulus frequency was 4.5 kHz for one *Triobp* mouse and 5 kHz for all other mice. Lines connect data from each mouse. (**e-f**) Absolute tonic OHC-DC displacements for 90 dB SPL stimuli as a function of age in *salsa* (**e**) and *Triobp* mice (**f**). Spearman’s rank correlations (ρ) and R^2^ values are provided in each panel, and dashed lines indicate linear fits. In **b**, **e**, and **f**, asterisks indicate statistical significance (* p < 0.05, ** p < 0.005, *** p < 0.0005).

In several WT and impaired mice, the dependence of the tonic displacements on OHC electromotility was tested by administering salicylate, which inhibits electromotility *in vitro* and *in vivo*^27–30^ (**Fig. 4c-d, Supplementary Fig. S4**). Tonic displacements were strongly reduced in all mice within 10-30 min of applying a small crystal of sodium salicylate to the round window membrane, where it was absorbed and transported apically. Vibrations at the stimulus frequency also sometimes changed after salicylate, possibly due to the reduction of a residual active component in the response or a slight drift in the measurement point. Regardless, these data demonstrate that the tonic displacements in both WT and impaired mice are attributable to OHC electromotility.

Though tonic displacements were observed in *salsa* and *Triobp* mice at a variety of ages, they tended to decrease in older mice (**Fig. 4e-f**). Responses were typically less than 5 nm and never more than 10 nm in mice 6 weeks of age or older. As the loss of tip links in *salsa* mice and overall deterioration of the stereociliary bundle in *Triobp* mice progress with age^24,25^, this trend could reflect a more complete loss of mechanotransduction currents in older mice. However, it is possible that the overall integrity of the OHCs also declines with age, leading to the reduction of any active OHC response. For instance, progressive OHC degeneration is commonly observed in mice with stereociliary mutations, including in *salsa* mice by 12 weeks of age^24^.

### Residual, nonlinear cycle-by-cycle OHC electromotility is observed in some impaired mice but is weakly correlated with tonic displacements

The lack of nonlinear BM amplification in *salsa* and *Triobp* mice suggests that cycle-by-cycle OHC electromotility is weak or absent. Nevertheless, electromotility more strongly influences the motion of the OHC region, which exhibits nonlinearity that is typically greater and more broadband than that observed for the BM^9,31,32^. Additionally, nonlinearity in OHC region motion can persist at low frequencies after application of drugs that eliminate BM nonlinearity^33,34^. Independent of the degree of nonlinearity, OHC contraction and elongation at the frequency of the stimulus should produce out-of-phase motions of the top and bottom of the OHC region^9^. To assess whether tonic OHC length changes in impaired mice may be accompanied by some degree of residual, cycle-by-cycle OHC electromotility, displacements of the OHC-DC junction and the TM were examined as a function of stimulus frequency and level. Responses were obtained from the TM instead of the RL, as the reflectivity of the latter was less stable, resulting in greater noise in the displacement measurements. In WT mice, the TM moves similarly to the RL^9^.

Nonlinearity in OHC region responses in WT mice was greatest at the CF but was also prominent at much lower frequencies (**Fig. 5a**). Average displacements of the OHC-DC junction and TM revealed nonlinear gain down to at least 2 kHz, one octave lower than the extent of the nonlinearity observed in BM displacements. The OHC-DC junction and TM also moved ∼0.5 cycles out of phase over a wide frequency range, consistent with cycle-by-cycle OHC elongation and contraction (**Fig. 5b**). At higher stimulus levels, the frequency extent of this out-of-phase motion was reduced, likely due to the increasing influence of the organ of Corti’s passive motion. However, even for 90 dB SPL stimuli, the TM and OHC-DC junction moved in antiphase at frequencies up to ∼3 kHz.

**Figure 5.**
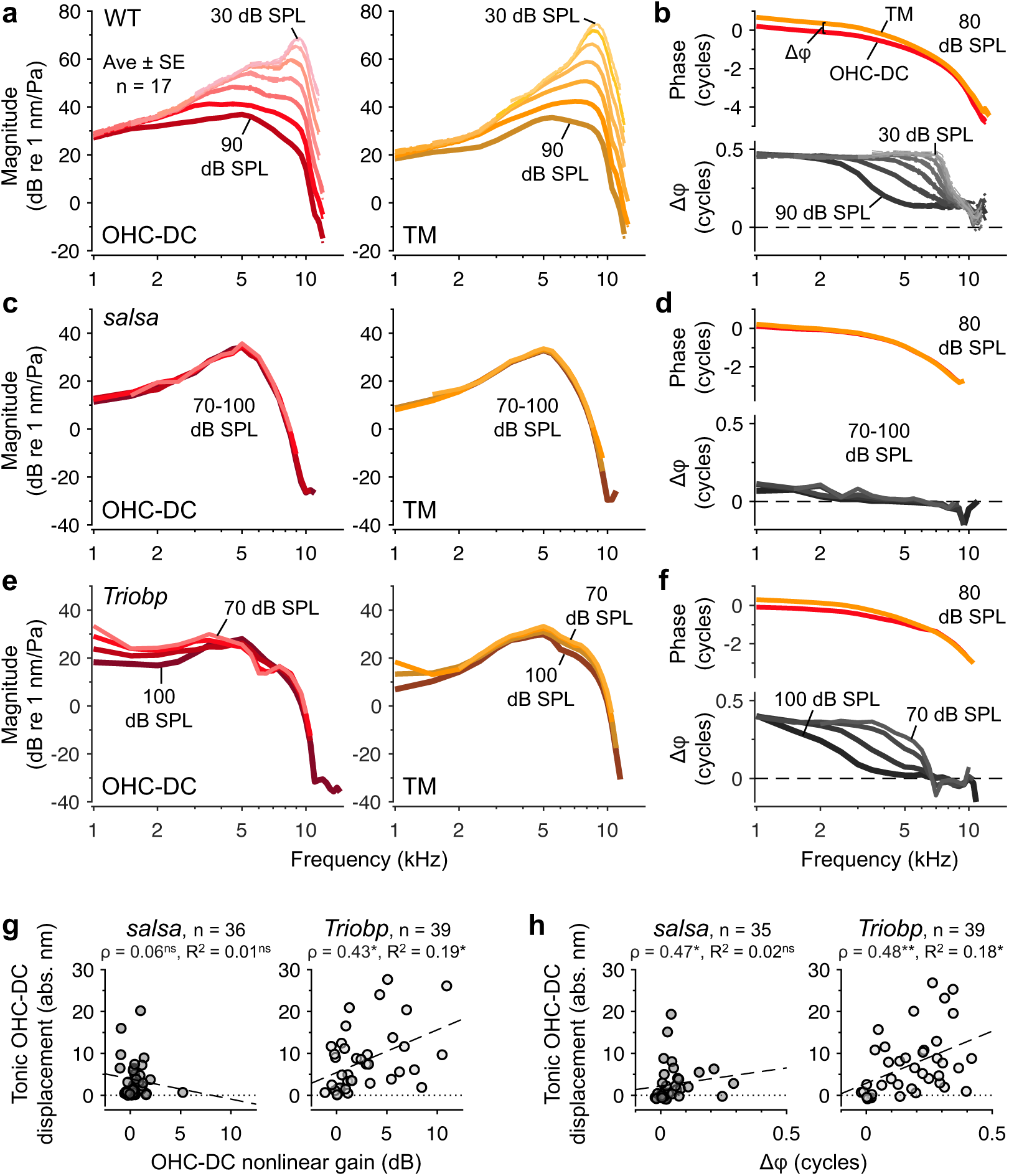
Nonlinear, cycle-by-cycle OHC electromotility is present in some impaired mice but is not strongly correlated with tonic displacements. (**a-b**) Average displacement gains (**a**) and phases (**b**) for the OHC-DC junction and TM as a function of frequency and stimulus level in WT mice (n = 17). Average phase differences between the TM and OHC-DC junction are shown in the bottom panel of **b**. Gains and phase differences are shown for stimulus levels of 30 to 90 dB SPL in 10 dB steps, while phases are shown only for 80 dB SPL stimuli, for clarity. Responses to higher levels are shown with darker/thicker lines. Dashed-dotted lines indicate ± 1 SE. Nonlinear gain was observed down to at least 2 kHz, and roughly out-of-phase motion was observed at low frequencies, even at high stimulus levels. (**c-d**) As in **a-b**, but for an individual *salsa* mouse. Gains and phase differences are shown for stimulus levels of 70 to 100 dB SPL in 10 dB steps, and phases are shown only for 80 dB SPL stimuli. Gain curves largely overlap, indicating response linearity, and phase differences are close to 0. (**e-f**) As in **c-d**, but for an individual *Triobp* mouse whose responses exhibited a small amount of nonlinearity and modest phase differences. Such nonlinearity and phase differences disappeared postmortem, as shown in **Supplementary Fig. S5**. (**g**) Absolute tonic OHC-DC displacement magnitudes plotted vs. the nonlinear gain observed in OHC-DC displacements for all *salsa* and *Triobp* mice. Nonlinear gain was quantified by taking the average difference between displacement gains at 70 and 100 dB SPL for stimulus frequencies of 1 to 4 kHz. (**h**) Absolute tonic OHC-DC displacement magnitudes plotted vs. the phase difference between TM and OHC-DC displacements for all *salsa* and *Triobp* mice. Phase differences were calculated for 80 dB SPL stimuli and averaged for stimulus frequencies of 1 to 4 kHz. In **g-h**, Spearman’s rank correlations (ρ) and R^2^ values are provided, with asterisks indicating significant correlations (* p < 0.05, ** p < 0.005, *** p < 0.0005, ns = not significant). Dashed lines are linear fits. Similarly weak-to-modest correlations were found when comparing nonlinearity observed in TM motion to tonic RL displacements, as shown in **Supplementary Fig. S6**.

In many impaired mice, OHC-DC and TM displacements lacked nonlinearity and phase differences at any frequency (e.g., **Fig. 5c-d**), indicating the absence of cycle-by-cycle electromotility. However, a small amount of nonlinearity along with modest phase differences could be observed in others, particularly in *Triobp* mice. Data from an individual *Triobp* mouse in **Fig. 5e-f** demonstrate that the nonlinearity in OHC-DC displacements was typically most prominent at lower frequencies, where responses for 70 dB SPL stimuli could exhibit up to ∼15 dB more gain than those for 100 dB SPL stimuli. Nonlinearity in TM responses was often observed across a broader frequency range, sometimes being strongest above 5 kHz. Roughly out-of-phase motion of the TM and OHC-DC junction was also observed at low frequencies in the example shown, with larger phase differences observed for lower stimulus levels (**Fig. 5f**). Phase differences were typically smaller than those in WT mice, presumably due to the weaker influence of the electromotile response relative to the passive organ of Corti motion.

When phase differences were detectable (i.e., greater than ∼0.1 cycle), low-frequency OHC-DC and TM motion generally lagged and led BM motion, respectively, following the same pattern observed in WT mice. In such cases, the average OHC-DC re BM phase (± SE) for a 2 kHz, 80 dB SPL stimulus was -0.18 ± 0.02 cycles, -0.12 ± 0.02 cycles, and -0.16 ± 0.02 cycles in WT (n = 17), *salsa* (n = 5), and *Triobp* (n = 25) mice, respectively, while the average TM re BM phase was 0.28 ± 0.02 cycles, 0.12 ± 0.03 cycles, and 0.13 ± 0.01 cycles. The phase relationship between OHC motility and BM motion was therefore similar to that in WT mice, suggesting that the phase of OHC stimulation was not dramatically altered in impaired mice. In all mice, any nonlinearity or phase differences largely disappeared postmortem (**Supplementary Fig. S5**).

Though the degree of nonlinearity in OHC-DC or TM responses could indicate the strength of residual mechanotransduction currents, this nonlinearity was poorly correlated with tonic displacement magnitudes in impaired mice (**Fig. 5g-h**, **Supplementary Fig. S6**). Nonlinearity in OHC-DC responses (quantified as the difference in gain for responses to 70 and 100 dB SPL stimuli, averaged from 1 to 4 kHz) was weakly but significantly correlated with tonic OHC-DC displacement magnitudes in *Triobp* mice, while no such trend was found in *salsa* mice, for which any nonlinearity was typically small (**Fig. 5g**). Nonlinearity in TM responses (quantified as the difference in gain for responses to 70 and 100 dB SPL stimuli, averaged from 1 to 9 kHz) was, however, significantly correlated with tonic RL displacements in both *salsa* and *Triobp* mice, though the correlations were weak to modest (**Supplementary Fig. S6**). The relatively weak correlations were not attributable to the choice of analysis parameters, as level-dependent gain differences exceeding 5 dB were observed in mice with little to no tonic displacement, and tonic displacements larger than 5 nm were found in mice lacking any discernible nonlinearity. For example, both the individual *salsa* and *Triobp* mice whose data are shown in **Fig. 5c-f** had measurable tonic displacements (with magnitudes of ∼7 and 19 nm, respectively; waveforms are shown in **Supplementary Fig. S4b** and **Fig. 4c**).

### Distortion-product otoacoustic emissions are present in some impaired mice but are also not strongly correlated with tonic displacements

Though intracochlear vibration measurements should most directly reveal any residual OHC electromotility, such activity could be missed due to the choice of measurement points or optical angle in any given mouse. Thus, distortion-product otoacoustic emissions (DPOAEs) were recorded in a subset of mice as a secondary assay of OHC function. Elicited by two stimulus tones at frequencies *f*_1_ and *f*_2_ (*f*_2_ > *f*_1_), DPOAEs are signals emitted to the ear canal at related frequencies like *2f*_1_-*f*_2_. The distortion is thought to be introduced by the nonlinear mechanotransduction process and transduced into electromotile force, with some of the resulting vibratory energy propagating out to the ear canal as a DPOAE.

While 2*f*_1_-*f*_2_ DPOAEs were measurable in some *salsa* and *Triobp* mice, they were typically present only at high stimulus levels (>75 dB SPL) and were much smaller than those measured in WT mice (**Fig. 6a**). For 85 dB SPL stimuli, DPOAEs were measurable above the background noise in ∼46% of *salsa* mice and ∼87% of *Triobp* mice tested. However, like the more direct measures of OHC nonlinearity and electromotility, DPOAE amplitudes were not strongly related to the tonic displacement magnitudes observed in impaired mice, though the correlations were statistically significant (**Fig. 6b**). When present, DPOAEs in impaired mice were reduced if not eliminated after death, demonstrating their physiological origin (**Supplementary Fig. S7**). Though the asymmetry that produces a tonic component should theoretically be more closely related to even-order distortions like *f*_2_-*f*_1_, *f*_2_-*f*_1_ DPOAEs were small in WT mice (less than 10 dB SPL for 85 dB SPL stimuli presented below 10 kHz), and were distinguishable from background noise and system artifact in only a handful of impaired mice.

**Figure 6.**
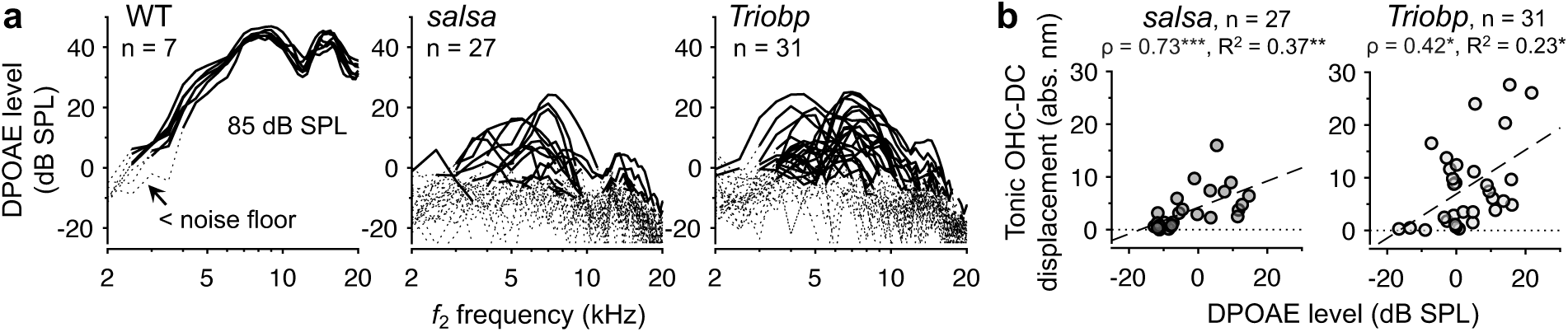
DPOAEs are present at high stimulus levels in some impaired mice and are modestly correlated with tonic displacement magnitudes. (**a**) DPOAE amplitudes in WT, *salsa*, and *Triobp* mice for 85 dB SPL stimuli plotted as a function of the *f*_2_ stimulus frequency (*f*_2_/*f*_1_ = 1.22). Data for all individual mice are overlaid and dotted portions of the curves indicate data not meeting the signal-to-noise criterion. (**b**) Absolute tonic OHC-DC displacement magnitudes plotted vs. DPOAE amplitudes for all *salsa* and *Triobp* mice. For each mouse, DPOAE amplitudes were averaged across *f*_2_ frequencies falling within ± 1 kHz of the frequency that elicited maximum BM displacement. Spearman’s rank correlations (ρ) and R^2^ values are provided, with significant correlations indicated by asterisks (* p < 0.05, ** p < 0.005, *** p < 0.0005). Dashed lines are linear fits. The physiological origin of the measured DPOAEs was confirmed by postmortem comparisons, as shown in **Supplementary Fig. S7**.

### Changes in the mechanotransducer function can produce tonic displacements in the absence of significant cycle-by-cycle amplification

The weak correlation between tonic displacement magnitudes and evidence of nonlinear, cycle-by-cycle OHC activity may suggest a dissociation between the mechanisms driving phasic and tonic motility.

However, using a simple OHC model with a Boltzmann function representing the mechanotransducer function, it was possible to reproduce the relative magnitudes of the tonic and phasic displacements both in WT mice and in impaired mice, after assuming realistic changes to the mechanotransducer function (**Fig. 7**).

**Figure 7.**
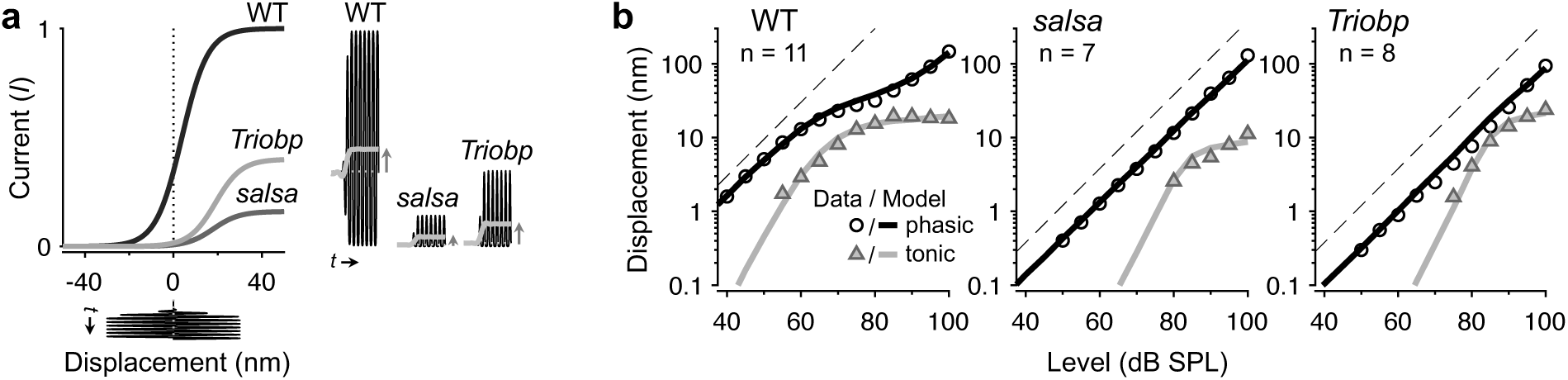
Changes in the mechanotransducer function can account for the presence of tonic responses in the absence of significant nonlinear amplification. (**a**) Boltzmann functions used to represent the relationship between OHC stereociliary bundle displacement and mechanotransduction current in WT, *salsa*, and *Triobp* mice. Waveforms are shown for a 5 kHz, 30 nm input and the corresponding outputs for each strain. Tonic components in the output waveforms are shown with gray curves. For all mice, the Boltzmann slope parameter *a* was 0.17 nm^-1^. In WT, *salsa*, and *Triobp* mice, respectively, the maximum current (*I*_max_) was 1, 0.16, and 0.4 and the operating point (*x*_0_) was 3.8, 18, and 19 nm. Active OHC region displacements were approximated by low-pass filtering the Boltzmann output and scaling by a factor of 125. To model the total OHC region displacement, the active response waveforms were summed with an estimate of the passive displacement, which was taken as the underlying BM displacement (i.e., the Boltzmann’s input) scaled by a factor of 0.56. (**b**) Average phasic and tonic displacements of the OHC-DC junction for 5 kHz tones varied in level (symbols) compared with modeled displacements (lines) in WT, *salsa*, and *Triobp* mice. Averages only include data meeting the signal-to-noise criterion from mice with tonic responses greater than 1.5 nm at 90 dB SPL and with BM displacements obtained in 5 dB steps over the same range of stimulus levels. Averages are shown when such clean data were obtained in at least 60% of mice. Responses from all individual mice, including those not meeting the above criteria, are shown in **Supplementary Fig. S8**. Dashed lines indicate linear growth.

In the model, OHC displacements were the vector sum of the cells’ passive displacement and their active electromotile response. The former was taken as BM displacement scaled by a factor of 0.56, which accounted for the translation of BM to OHC region motion observed postmortem. The latter was approximated by the low-pass-filtered output of a first-order Boltzmann function, which has been shown to replicate the relative magnitudes of tonic, harmonic, and intermodulation distortions in WT mice^9,35^. The Boltzmann function represents the relationship between stereocilia displacement (*x*) and mechanotransducer current (*I*), and is given by:

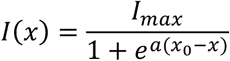

where *I*_max_ is the maximum output current, *a* is a slope parameter (in nm^-1^), and *x*_0_ is the resting position, or operating point, of the function (in nm).

Using average BM displacements as a proxy for stereocilia displacement, the resulting output was filtered with a first-order, low-pass filter (corner frequency of 1.75 kHz) to account for shaping of the receptor potential by the OHC membrane’s electrical properties^6,9,21^. The active displacement response was then assumed to be proportional to the low-pass filtered Boltzmann output. Due to the simplifications above, the output of the Boltzmann was unitless, and *I*_max_ was set to 1 when modeling WT responses. The active component was therefore scaled prior to summing with the passive displacement (by a factor of 125, as determined through fitting).

In WT mice, the Boltzmann parameters and scaling factor were adjusted so that the growth and relative magnitudes of the phasic and tonic displacements in the modeled OHC motion matched those in average OHC-DC junction displacements for 5 kHz stimuli varied in level (**Fig. 7a-b**). The fitted Boltzmann parameters (*x*_0_ = 3.8, *a* = 0.17) were similar to those previously estimated in CBA/CaJ mice for responses to CF tones^9^, and corresponded to ∼34% of the mechanotransduction current being activated at rest.

In *salsa* and *Triobp* mice, it was also possible to approximate the phasic and tonic components of average OHC region displacements to 5 kHz tones by simply reducing the maximum output current of the Boltzmann function and positioning the operating point away from the center of the function, along its left-most saturating edge (**Fig. 7a**). Both of these changes dramatically reduced the phasic response in the Boltzmann’s output, thus minimizing the active component of OHC motion at the stimulus frequency. As a result, the total phasic displacement was dominated by the passive motion and grew linearly with stimulus level (**Fig. 7b**). However, due to the shifted operating point, the Boltzmann output was strongly asymmetric, leading to the presence of a significant tonic component in the active response. Depending on the parameters used, the tonic response could grow to become as large or larger than that observed in WT mice for high stimulus levels. This simple model therefore demonstrates how residual, asymmetric mechanotransduction currents can lead to the observation of active, and sometimes quite large, tonic OHC contractions, despite apparently linear and passive motion at the stimulus frequency.

It is reasonable to assume that the maximum current and operating point are altered in the impaired mice. In *salsa* mice, loss of tip links would decrease the total number of functional mechanotransduction channels, strongly reducing the maximum current^24^. The loss of tension among stereocilia would also lead to channel closure, including the closure of channels that may still be gated by any remaining tip links.

This could shift the bundle’s operating point toward a position where the resting current was near 0%. In *Triobp* mice, similar effects may result from overall bundle degeneration, though the presence of tip links would lead to a greater number of functional channels than in *salsa* mice, resulting in a larger maximum current (estimated to be 40% of the WT value, compared to 16% of the WT value in *salsa* mice).

Importantly, the heterogeneity in the observed responses (see **Supplementary Fig. S8**) can be accounted for by manipulating the Boltzmann parameters. Phasic responses with mild degrees of compressive nonlinearity are achieved when the maximum current is set sufficiently high, tonic responses of opposite polarity are produced by shifting the operating point to the right-most saturating edge, leading to primarily hyperpolarizing currents, and no tonic responses are produced when the operating point is centered. Thus, variations in the mechanotransducer function across mice can explain the diversity of responses, as well as the weak correlations between tonic displacements and signs of residual phasic electromotility.

## DISCUSSION

The present data show that sound-evoked tonic OHC motility can persist in mice lacking normal mechanotransduction and cochlear amplification. Tonic OHC responses are therefore not specific indicators of sensitive cochlear function. While the data could suggest that distinct mechanisms underlie tonic and phasic OHC motility, the relationships between tonic and phasic motions in both WT and impaired mice were successfully reproduced using a model in which all nonlinearity originated from the mechanotransducer function. Nevertheless, it is important to consider mechanisms other than conventional mechanotransduction that may contribute to or even dominate the generation of the tonic OHC responses observed here.

In the absence of conventional mechanotransduction (i.e., that mediated by channels located near the tips of the stereocilia), sound-induced deformation of the OHC membrane may elicit current flow through channels located along the apical or basolateral aspects of the cell membrane^36–38^. In particular, PIEZO2 channels located on the apical OHC surface have recently been shown to produce stretch-activated currents that are revealed when conventional mechanotransduction is silenced by genetic manipulations, tip link breakage, or general damage to the bundle^39^. PIEZO2 channels may therefore offer an alternative current path in *salsa* and *Triobp* mice. However, it remains unclear whether PIEZO2 or other alternative channels are activated by sound-evoked motion of the organ of Corti *in vivo*, or if stimulation at kilohertz rates can elicit substantial tonic currents. It is also uncertain whether eliminating PIEZO2 in impaired mice would sufficiently distinguish between the role of conventional vs. non-conventional mechanotransduction in eliciting tonic OHC motility. PIEZO2 has been suggested to facilitate high-frequency mechanotransduction^40^ and even form part of the conventional mechanotransduction machinery^41^. Furthermore, conditional knockout of PIEZO2 channels leads to varying degrees of hearing loss^39–41^. The roles of apical and basolateral conductances in both normal and impaired OHC function therefore require further study.

Possibly also relevant to the present observations is the finding that mechanically stimulating the somas of isolated guinea pig OHCs can elicit both tonic and phasic motility^11,12,42^. The resulting tonic motions can exceed the phasic responses and approach 100 nm at high stimulus intensities^42^. However, it is unclear if this mechanically induced motility involves the same processes that underlie electromotility, if it depends on the stretch-activated conductances discussed above, or if it stems from general piezoelectric effects^43–45^. If tonic responses in impaired mice were due to mechanically induced motility, the near elimination of the responses postmortem suggests either that this form of motility is not activated by the strain caused by passive organ of Corti vibrations, or that it is sensitive to factors influenced by death, such as the endocochlear potential or OHC turgor pressure. The contribution of mechanically induced motility to responses in WT mice likewise remains obscure.

It is important to consider that the electromotility of isolated OHCs is inherently asymmetric, such that the tonic OHC motility observed *in vivo* may not solely depend on tonic current flow. Specifically, for OHCs held at near-physiological resting potentials *in vitro*, depolarization-induced contractions saturate at magnitudes that are larger than those of hyperpolarization-induced elongations^46,47^. When driven by a sinusoidally varying voltage, OHCs would therefore be observed to tonically contract. However, such asymmetry may only be appreciable when examined over relatively large changes in membrane potential. If sound-evoked receptor potentials do not exceed ∼10 mV^22,48^, the response asymmetry contributed by electromotility is expected to be small^49^. Furthermore, an asymmetry in the electromotile response alone would not produce tonic displacements that exceed the phasic displacements^9,17^.

Various mechanisms may also shape mechanotransduction currents and motile responses in impaired mice differently than in WT mice, perhaps rendering them more low-pass in nature. Bundle stiffness is reduced in *Triobp* (and presumably, *salsa*) mice^25^, and the overall bundle disorganization may further act to favor low-frequency motion and/or current flow. Disruptions in the resting OHC conductance (e.g., due to channel closure), may also lower the corner frequency of the membrane filter that shapes the receptor potential^21^, causing greater low-pass filtering of the drive to electromotility. Lastly, the larger tonic responses in *Triobp* compared to *salsa* mice could reflect differences in the passive mechanical properties of the OHCs and organ of Corti. For instance, atomic force microscopy has revealed that the RL in *Triobp* mice is less stiff than in WT and has an altered stiffness gradient along its width^50^. This could result in larger deformations of the RL and surrounding regions in response to a tonic OHC length change.

Regardless of the underlying mechanisms, it is unlikely that tonic OHC responses play a significant mechanical role in impaired mice. However, if these motile responses are driven by residual mechanotransduction currents, their presence may indicate that the OHCs remain viable and could be targeted for functional recovery. Alternatively, they may simply be an epiphenomenon of a purely aberrant, pathological state. Whether tonic responses in impaired and normal mice are produced by the same mechanisms, and whether they play a role in modulating cochlear amplification under normal conditions, remains to be more fully explored.

## METHODS

### Mice

Measurements were obtained from 3-7 week-old mice with roughly equal numbers of male and female mice used for each strain. *salsa* mice were originally generated at Scripps Research Institute by Dr. Ulrich Mueller, while *Triobp* mice were generated at the NIH/NIDCD by Dr. Thomas Friedman. Both *salsa* and *Triobp* mice were provided to Dr. John Oghalai at Stanford University and eventually re-derived by Charles River before being re-established at the University of Southern California, where they have been housed since 2017. WT C57BL/6J mice were obtained from Jackson Laboratories and then bred on-site. All procedures were approved by the local Institutional Animal Care and Use Committee at the University of Southern California.

A total of 18 WT (10 female), 46 *salsa* (24 female), and 44 *Triobp* (22 female) mice were used over the course of this study. Data collection was not attempted in some of these mice due to premature death during surgery (for three *salsa* mice and one *Triobp* mouse), the presence of fluid or abnormal tissue in the middle ear space (for three salsa and four *Triobp* mice), or calibration-related issues (for one *Triobp* mouse). In each experiment, mice were anesthetized (80-100 mg/kg ketamine, 5-10 mg/kg xylazine) and placed on a heating pad set to 38 °C. Supplemental doses of anesthesia were provided throughout the experiment to maintain areflexia. After fixing the exposed skull to a head-holder with dental cement, the left bulla was surgically accessed. The bone below the tympanic annulus was carefully removed to reveal the middle ear space, providing optical access to the apical cochlear surface. The pinna and portion of the ear canal were resected so that the tip of an acoustic probe (ER10-X; Etymotic Research, Inc., Elk Grove Village, IL) could be placed within 3 mm of the eardrum. The probe was coupled to the residual ear canal with plastic tubing, which was sealed in place with dental cement. A tracheotomy was often performed to facilitate free breathing. All experiments were conducted on a vibration-isolating table in a sound-attenuating booth.

### OCT imaging and vibrometry

Sound-evoked cochlear vibrations were measured using a custom-built OCT system that has been described previously^9,51–53^. The system used a swept light source with a 1310 nm center wavelength and 95 nm bandwidth (Insight Photonic Solutions; Lafayette, CO). The axial and lateral imaging resolutions were 12.5 and 9.8 microns, respectively.

The light source was first scanned across the cochlear bone to obtain cross-sectional images of an apical location tuned to ∼9 kHz in WT mice. Individual pixels in the image were then chosen for vibration measurements. The angle of the BM relative to the optical path was between 60 and 90°, such that the measurements primarily captured the transverse motion of the different structures. Nevertheless, the measurements likely also included some component of radial and/or longitudinal motion. Measurements were made from points that were particularly bright and stable, as these yield data with higher signal-to-noise ratios and reduced artifact^54^. Since the reflectivity of individual points could change over time, cross-sectional images were regularly examined throughout each experiment. Vibratory responses were obtained with a sampling rate of 100 kHz. All acoustic stimuli were calibrated using the pressure measured by the probe microphone *in situ*.

Displacements were first obtained for tones varied in frequency (1-15 kHz in 0.5 kHz steps) and sound pressure level (SPL; 10-90 dB SPL in WT mice; 50-100 dB SPL in impaired mice), so as to characterize their tuning and nonlinearity. Measurements were typically obtained from the BM, TM, and OHC-DC junction. The growth of responses at specific frequencies was also sometimes characterized using finer level steps (3 or 5 dB). Frequency and level functions were obtained with 102 ms tones (including 1 ms onset and offset ramps) presented once every 110 ms. To obtain the amplitude and phase of the displacement at the stimulus frequency, the steady-state portion of the average response to 4-8 stimulus repetitions was analyzed via fast Fourier transform (FFT). The mean + 3 standard deviations (SDs) of the displacement magnitudes 140-240 Hz below and above the stimulus frequency were taken as the measurement noise floor.

After obtaining frequency responses, tonic displacements of various points were then assessed for 90 dB SPL stimuli presented at a frequency within 0.5 kHz of that eliciting the largest BM displacement. This frequency was also found to elicit the largest tonic displacements. In impaired mice, this frequency was individually determined for each mouse and fell between 4 and 5.5 kHz. For WT mice, responses were always obtained at 5 kHz in order to facilitate averaging across mice. Responses at other frequencies and levels were also sometimes obtained when the reflectivity of a given measurement point was sufficiently stable.

Tonic displacements were typically measured in response to 7 ms tones (including 1 ms onset and offset ramps) presented once every 20 ms. In the first three *salsa* mice tested, 12 ms tones were used. Responses were obtained for 500-6,000 stimulus repetitions, depending on stimulus level and the reflectivity of the measurement point. A large number of repetitions was needed to reduce the influence of low-frequency motion artifacts. To further minimize such artifacts, final response averages were computed from the mean of 50 sub-averages. Each sub-average included 40% of the responses, selected randomly and with replacement, and was retained in the final average if the pre- and post-stimulus intervals in the waveform differed by less than 1.5 nm. Pre- and post-stimulus intervals were defined as the 2 ms immediately prior to stimulus onset and the 3-11 ms window after the stimulus offset, respectively.

To visualize the tonic displacements, average waveforms were low-pass filtered to eliminate any phasic response at the stimulus frequency. The corner frequency of the first-order filter was 0.3 kHz for 1 kHz stimuli, 0.5 kHz for 2-3 kHz stimuli, and 1 kHz for higher frequencies. Tonic displacement magnitudes were quantified by subtracting the average filtered waveform values in the 1 ms prior to stimulus onset from the average displacement occurring 4-5 ms post-stimulus onset. The measurement noise was estimated by taking the absolute difference between average filtered displacement values in the pre- and post-stimulus intervals and adding 3 SDs of the values compiled across both intervals. An FFT was also applied to the steady-state portion of the average unfiltered response waveforms to determine the amplitude and phase of the response at the stimulus frequency. For this purpose, the measurement noise floor was taken as the mean plus 3 SDs of the response magnitudes 0.6-1.6 kHz below and above the stimulus frequency.

The above analysis method captured the steady-state tonic response in order to avoid the effects of any distortions in the speaker output at the end of the stimulus onset ramp. Such distortions could be observed at high stimulus levels, though were generally small. Tonic displacements did occasionally adapt post-stimulus onset, even in the absence of such acoustic distortions, such that the values reported here may be lower than the maximum tonic displacement observed. However, the amount of adaptation was typically less than 20% in most measurements. Using the maximum instead of the steady-state tonic displacements did not meaningfully change the results.

Measurements were conducted for as long as a mouse’s condition remained stable, which was usually 2 to 6 hours. Mice sometimes succumbed to the effects of anesthesia and measurements were terminated upon signs of labored breathing or cardiac issues (e.g., reduced or arrhythmic heartbeat). If all desired measurements were completed prior to this point, mice were euthanized via anesthetic overdose. A subset of measurements was then repeated up to 60 min postmortem. At later postmortem times, the response magnitudes and tuning can change^26,55^, possibly indicating alterations of cochlear and/or middle ear function that confound interpretation of the measurements. Postmortem measurements typically included frequency responses at stimulus levels of 50 to 100 dB SPL in 10 dB steps and tonic responses for 90 dB SPL stimuli.

Preparations were sometimes compromised by fluid leaking into the middle ear space or deterioration of the OCT images, possibly due to optical changes resulting from drying of the cochlear bone. In such cases, or if all desired measurements were successfully completed, mice were euthanized and postmortem measurements were either not attempted or not included in any of the analyses presented here.

### Salicylate administration

In several live mice, a small crystal of sodium salicylate (Spectrum Chemical Mfg. Corp.; Gardena, CA) was applied to the round window membrane in order to inhibit OHC electromotility^27–30^. Postmortem responses from mice treated with salicylate were not included in any analyses.

### Distortion-product otoacoustic emissions (DPOAEs)

DPOAEs were measured in response to two stimulus tones (at frequencies *f*_1_ and *f*_2_, *f*_2_ > *f*_1_), with *f*_2_ ranging from 2-24 kHz in 0.5 kHz steps, and an *f*_2_/*f*_1_ ratio of 1.22. The levels of the stimuli were equal and typically varied from 60-85 dB SPL in 5 dB steps. Stimuli were 102 ms tones (including 1 ms onset and offset ramps) and presented once every ∼110 ms. In impaired mice, responses were averaged over 24 stimulus repetitions. In WT mice, responses were averaged over only 8 repetitions to minimize any fatiguing effects from exposure to the higher intensity stimuli. The magnitude and phase of the 2*f*_1_-*f*_2_ DPOAE were taken from the FFT applied to the steady-state portion of the average response waveform. Signals with magnitudes greater than the mean + 3 SDs of the pressure measured 40-140 Hz below and above the DPOAE frequency were considered to be above the measurement noise floor.

When DPOAEs were measurable in live mice, recordings were also obtained postmortem for ∼80-85 dB SPL stimuli to determine the possible presence of system artifacts. While postmortem DPOAEs were sometimes present at these high stimulus levels, they were almost always considerably reduced compared to the *in vivo* responses. In three WT mice, postmortem measurements repeated over ∼1.5 hours also revealed that DPOAEs continued to decrease over time, further suggesting that the responses were physiological. Only in one *salsa* mouse were postmortem responses similar in magnitude to those measured *in vivo*, though the DPOAEs were close to the measurement noise floor in either condition.

These data were excluded from subsequent analyses.

### Data reporting and analysis

All displacement magnitudes are peak values and phases are referenced to the phase of the stimulus pressure measured in the ear canal. Group data are reported as the mean ± 1 standard error (SE) unless otherwise specified. Individual and average displacement data plotted as a function of frequency or stimulus level only include data with magnitudes above the measurement noise floor. Average data are only shown when such data were available from at least 60% of the mice tested. In plots of displacement vs. stimulus frequency or level, isolated data points or segments were sometimes excluded for clarity. Statistical analyses were performed in SPSS (IBM Corp.; Armonk, NY) or MATLAB (MathWorks; Natick, MA) with p values < 0.05 indicating a significant result. Comparisons of response properties across mouse strains or across measurement locations within a given strain were conducted using ANOVA, with Bonferroni corrections applied to post-hoc analyses. Pre- and postmortem measurements were compared with two-sided, paired t-tests.

## ACKNOWLEDGMENTS

This work was supported by NIH/NIDCD R21 DC019209 and R01 R01DC021006, as well as the University of Southern California’s Keck School of Medicine. I thank Dr. Ulrich Mueller and Dr. Thomas Friedman for originally providing the *salsa* and *Triobp* mice, respectively.

## Author Contributions

J.B.D. designed and performed research, analyzed data, and wrote the paper.

