## Supplementary Material for "Tonic sound-evoked motility of cochlear outer hair cells in mice with impaired mechanotransduction"

James B. Dewey

This file contains:
Supplementary Figures S1 to S8
Supplementary References

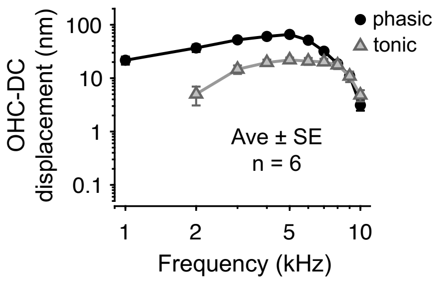


**Supplementary Figure S1. High-level stimuli elicit tonic displacements in WT mice that are broadly tuned and peak near 5 kHz.** Average displacements of the OHC-DC junction in six WT mice for 90 dB SPL tones varied from 1 to 10 kHz in 1 kHz steps. Tonic displacements were largest near ~5 kHz, where the phasic displacements of the OHC-DC junction and BM were also maximal (average BM response are shown in **Fig. 2d** of the main text). Averages only include data that were above the measurement noise floor and are plotted if such data were available in at least four mice. Error bars indicate ± 1 SE and are typically smaller than the symbols used to plot the average values.

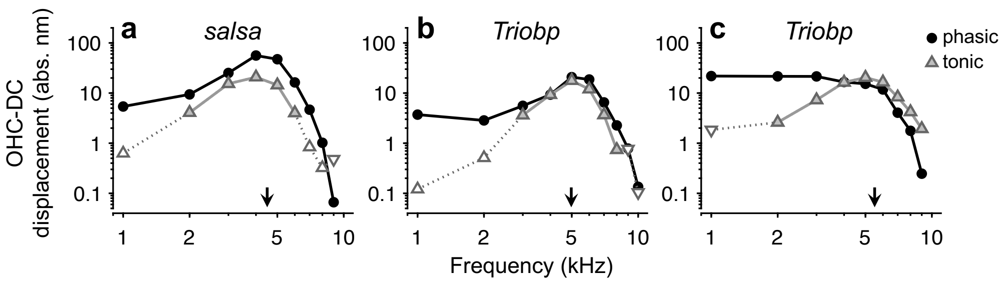


**Supplementary Figure S2. Tonic displacements in impaired mice also peak near 5 kHz.** (**a**-**c**) Phasic and absolute tonic displacements of the OHC-DC junction in one *salsa* mouse (**a**) and two *Triobp* mice (**b**-**c**). Responses were obtained for 90 dB SPL tones varied from 1 to 9 or 10 kHz in 1 kHz steps. Tonic displacements peaked near the frequency eliciting maximal BM displacement in each mouse (indicated by arrows in each panel). This was not always the same frequency as that eliciting maximal phasic OHC-DC displacements, which occasionally had a more low-pass characteristic, as in **c** (frequency responses of the OHC-DC junction are examined in greater detail in **Fig. 5** of the main text). In each panel, dotted lines and open triangles indicate frequencies where tonic displacements were not distinguishable from measurement noise. Such displacement values were occasionally negative (i.e., toward scala tympani), in which case they are shown with downward pointing triangles. For the examples provided here, all tonic displacements that were confidently distinguishable from noise were in the direction of scala vestibuli and are shown with upward pointing triangles.

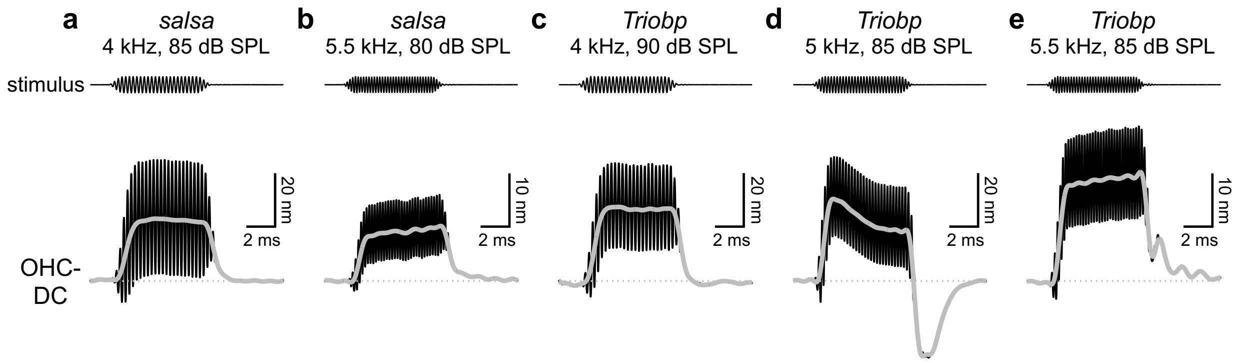


**Supplementary Figure S3. Tonic displacements can exceed phasic displacements in impaired mice.** (**a**-**e**) OHC-DC junction displacement waveforms from two *salsa* mice (**a**-**b**) and three *Triobp* mice (**c**-**d**). The stimulus frequency and level for each measurement are indicated above each stimulus waveform. Low-pass-filtered displacement waveforms (gray curves) reveal the tonic displacements, which are larger than the phasic displacements in all examples shown. The individual waveforms shown here were also chosen to highlight the diversity in the morphology of the tonic responses, which could increase or decrease over time. At the offset of the stimulus, some responses also exhibit a tonic shift toward positions beyond the initial pre-stimulus baseline, followed by gradual adaptation toward the baseline over several ms (as in **d**). The origins of these behaviors are unclear, but could involve opening and closing of voltage-gated channels in the basolateral membrane[1–3], or stereociliary bundle adaptation[4–6]. Low-frequency oscillations were also observed after the stimulus offset in several mice (as in **e**). Note that displacement scale bars indicate either 10 or 20 nm.

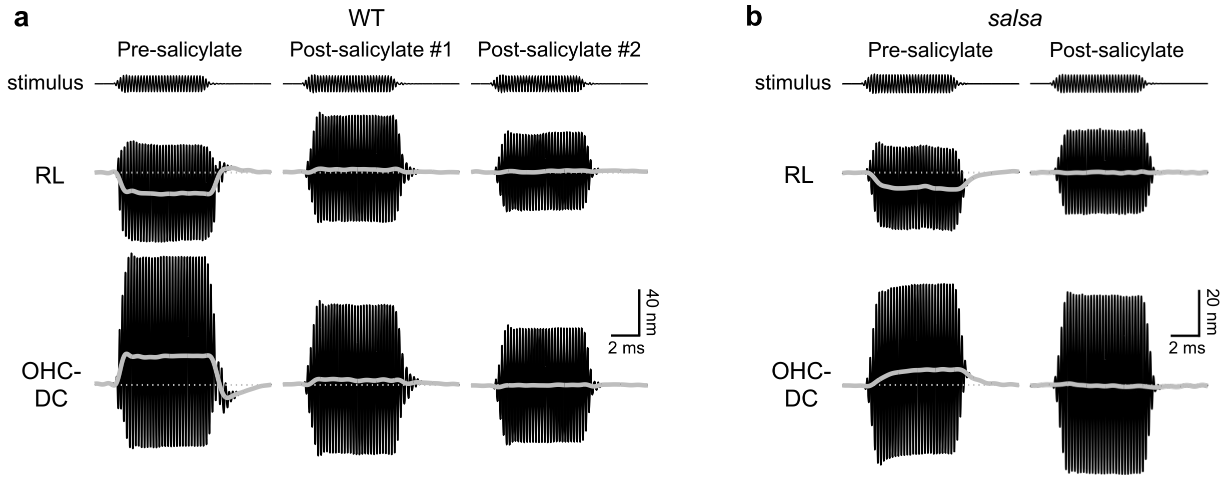


**Supplementary Figure S4. Salicylate reduces tonic OHC motility in WT and impaired mice.** (**a**) Waveforms of RL and OHC-DC junction displacements elicited by a 5 kHz, 90 dB SPL tone in a WT mouse before and after one or two applications of salicylate to the round window membrane. Tonic displacements were dramatically reduced within 10 min of the first salicylate application (as shown by waveforms under 'Post-salicylate #1'). However, small tonic displacements of both the RL and OHC-DC junction toward scala vestibuli persisted. A second application of salicylate ~1 hr after the first more completely abolished the tonic displacements (recordings shown under 'Post-Stimulus #2' were obtained ~30 min after the second application). Partial recovery was observed at later time points, though the recovery process was not systematically tracked. (**b**) Pre- and post-salicylate displacement waveforms for the RL and OHC-DC junction in a *salsa* mouse. A single application of salicylate was sufficient to eliminate the tonic displacements. Pre- and post-salicylate tonic displacements are quantified in **Fig. 3d** of the main text, with post-salicylate data obtained after one application of salicylate for all mice. Note that different displacement scales are used in **a** and **b**. Stimulus waveforms are arbitrarily scaled.

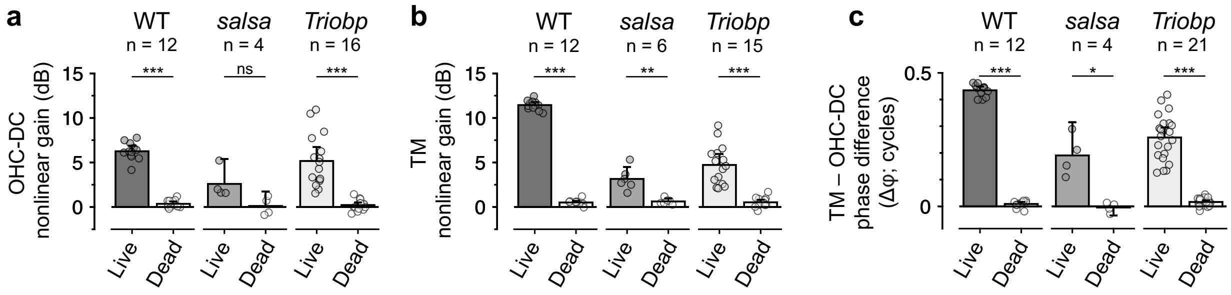


**Supplementary Figure S5. Nonlinearity and phase differences in WT and impaired mice are strongly reduced postmortem.** (**a**) Comparison of the nonlinear gain observed in OHC-DC displacements pre- and postmortem in all mice that exhibited at least 1.5 dB of nonlinear gain before death. Bar graphs show average values (with error bars indicating 95% confidence intervals) and individual data are shown with semi-transparent symbols. Nonlinear gain was quantified by taking the difference between displacement gains for responses to 70 and 90 dB SPL stimuli in WT mice, and 70 and 100 dB SPL stimuli in *salsa* and *Triobp* mice, and then averaging such gain differences from 1 to 4 kHz. Nonlinearity decreased after death in all mice, though the comparisons of pre- and postmortem values with paired t-tests were only significant for WT and *Triobp* mice (for WT, $t_{11} = 19.00$, $p < 0.0005$; for *salsa*, $t_4 = 2.01$, $p = 0.14$; for *Triobp*, $t_{16} = 6.41$, $p < 0.0005$). Postmortem changes were not significant in *salsa* mice due to the low number of live mice with at least 1.5 dB of nonlinear gain. (**b**) As in **a**, but for nonlinear gain observed in TM displacements, with gain differences averaged from 1 to 9 kHz. A wide frequency range was used to assess TM nonlinearity, as gain differences were often observed more uniformly across frequency when compared to those observed for the OHC-DC junction. Nonlinearity was significantly decreased after death in all strains, as assessed by paired t-tests (for WT, $t_{11} = 76.66$, $p < 0.0005$; for *salsa*, $t_5 = 6.30$, $p = 0.001$; for *Triobp*, $t_{14} = 6.74$, $p < 0.0005$). (**c**) As in **a** and **b**, but for the comparison of phase differences between TM and OHC-DC junction motion observed pre- and postmortem in all mice that exhibited at least 0.1 cycles of phase difference before death. Phase differences were evaluated for 80 dB SPL stimuli and averaged from 1 to 4 kHz. Phase differences were significantly decreased after death in all strains, as assessed by paired t-tests (for WT, $t_{11} = 52.61$, $p < 0.0005$; for *salsa*, $t_3 = 4.17$, $p = 0.03$; for *Triobp*, $t_{20} = 12.33$, $p < 0.0005$). In **a-c**, asterisks indicate significant differences between pre- and postmortem values (* $p < 0.05$, ** $p < 0.005$, *** $p < 0.0005$, ns = not significant).

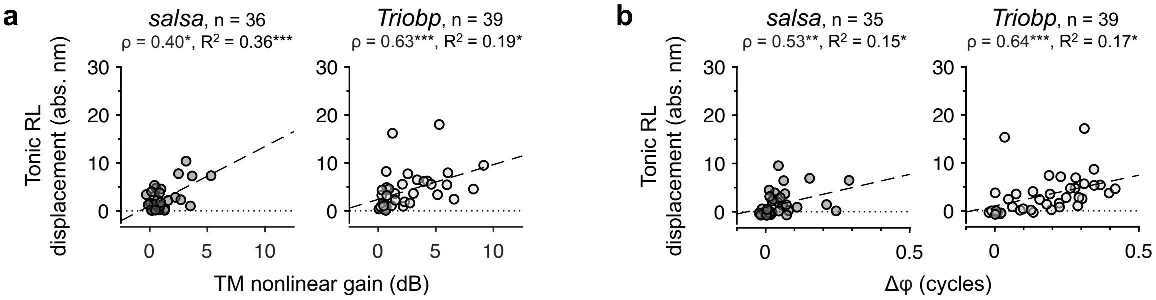


**Supplementary Figure S6. Tonic RL displacements are modestly correlated with nonlinearity in TM displacements and phase differences between TM and OHC-DC junction displacements in impaired mice.** (**a**) Absolute tonic RL displacement magnitudes plotted vs. the nonlinear gain observed in TM displacements for all *salsa* and *Triobp* mice. Nonlinear gain was quantified by taking the average difference between displacement gains at 70 and 100 dB SPL for stimulus frequencies of 1 to 9 kHz. (**b**) Absolute tonic RL displacement magnitudes plotted vs. the phase difference (Δφ) between displacements of the TM and OHC-DC junction for all *salsa* and *Triobp* mice. Phase differences were calculated for responses to 80 dB SPL stimuli and averaged over frequencies of 1 to 4 kHz. In **a** and **b**, Spearman's rho (ρ) and $R^2$ values are provided, with asterisks indicating significant correlations (* $p < 0.05$, ** $p < 0.005$, *** $p < 0.0005$). Dashed lines are linear fits.

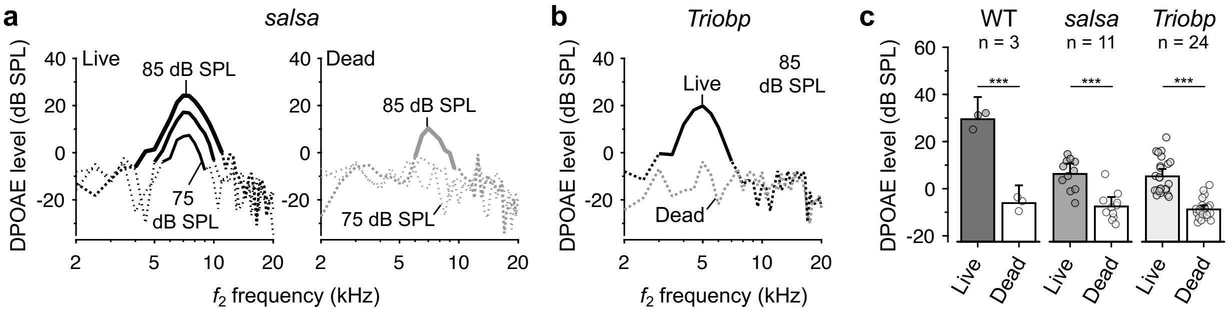


**Supplementary Figure S7. DPOAEs are reduced postmortem in WT and impaired mice.** (**a**) DPOAE amplitudes in an individual *salsa* mouse obtained pre- and postmortem. Responses are shown for 75, 80, and 85 dB SPL stimuli, with dotted portions of the curves indicating data not meeting the signal-to-noise criterion. While postmortem DPOAEs were sometimes measurable at high stimulus levels, they were always smaller than those obtained premortem. This indicates that the DPOAEs in live impaired mice were of physiological origin. (**b**) DPOAE amplitudes in an individual *Triobp* mouse obtained pre- and postmortem for 85 dB SPL stimuli. In this example, DPOAEs were not detectable above the measurement noise floor after death. (**c**) Comparison of pre- and postmortem DPOAE amplitudes for 85 dB SPL stimuli in WT, *salsa*, and *Triobp* mice. Bar graphs show average values (with error bars indicating 95% confidence intervals) and individual data are shown with semi-transparent symbols. For each mouse, DPOAE amplitudes were averaged over a 2 kHz range centered on the frequency eliciting maximum BM displacement (which fell between 4 and 5.5 kHz). Only data from mice in which the premortem DPOAE amplitude was at least -10 dB SPL and above the average measurement noise floor are included. DPOAE amplitudes were reduced postmortem in all strains, as assessed by paired t-test (for WT, $t_2 = 67.25$, $p < 0.0005$; for *salsa*, $t_{10} = 7.05$, $p < 0.0005$; for *Triobp*, $t_{23} = 10.75$, $p < 0.0005$). Asterisks indicate significant differences (*** $p < 0.0005$).

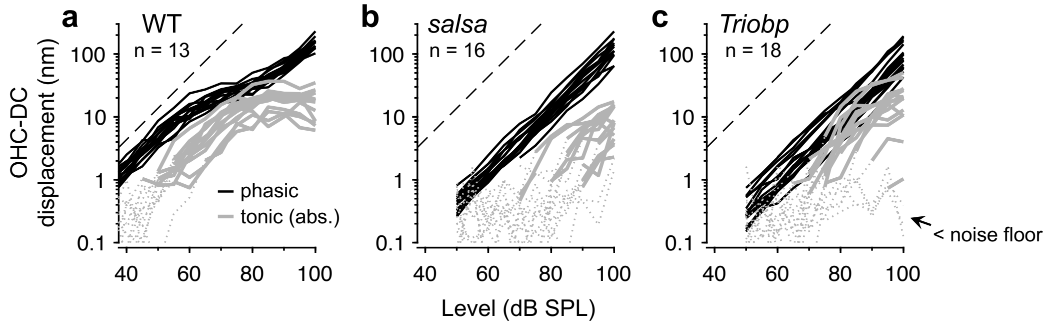


**Supplementary Figure S8. Growth of tonic and phasic displacements vs. stimulus level for 5 kHz tones in WT and impaired mice.** (**a**) Displacements of the OHC-DC junction in WT mice for 5 kHz tones varied in level in 5 dB steps. Phasic responses at the stimulus frequency (black curves) and absolute tonic displacements (gray curves) are shown overlaid for all mice. Dotted portions of the curves indicate data not meeting the signal-to-noise criterion. (**b-c**) As in **a** but for *salsa* (**b**) and *Triobp* mice (**c**). To best illustrate the heterogeneity in the responses, data are shown for all mice in which such measurements were obtained, including those in which the tonic responses were indistinguishable from background noise at all stimulus levels. The datasets shown here are therefore larger than those used to fit the Boltzmann-based model in **Fig. 7** of the main text. The latter only included data from mice with an absolute tonic response greater than 1.5 nm at 90 dB SPL and in which BM displacements were also obtained in 5 dB steps.
